# Loss of YAP/TAZ impaired the proliferation and differentiation ability of neural progenitor cells

**DOI:** 10.1101/296053

**Authors:** Shanshan Kong

**Affiliations:** University of Tennessee Health Science Center, 50 North Dunlap Street, Memphis, TN, 38103, USA

## Abstract

YAP (Yes-associated protein) and TAZ (transcriptional coactivator with PDZ-binding motif) are downstream effectors of the Hippo pathway, they activate the expression of transcriptional targets that promote cell growth, cell proliferation, and prevent apoptosis. Here I examined the function of YAP/TAZ in mouse neocortex development through conditional deletion of Yap and Taz by Emx1-Cre. Loss of YAP/TAZ cause the hydrocephalus after birth, leads to aberrant development and dilated ventricle in adult stage, this phenotype can be detected as early as P0. YAP/TAZ are expressed in Sox2+ neural progenitor cells, when YAP/TAZ are deleted, the neuroepithelial cell junctions are disrupted; the numbers of Sox2+ cell and Tbr2+ cell are reduced and the ratio of tbr2/Sox2 is also reduced at E15.5. Results of cell cycle analyzing experiments display YAP/TAZ deletion increased the cell cycle exit. The improperly increased expression of Tuj1+ in progenitor cells in the YAP/TAZ deleted cortex indicates the premature of Sox2+ progenitor cells. Together, our results reveal that YAP/TAZ deletion changed the polarity of neuroepithelial cells, and increased the cell cycle exit, reduced the differentiation of Sox2+ cells into Tbr2+ cells through promoting the premature of Tuj1+ cells. These results define the functions of YAP/TAZ in keeping the cell polarity neural progenitors and ensuring their proliferation and differentiation, and also reveal the roles of YAP/TAZ in developing cortex.

## Introduction

Hippo pathway is the one of the fundamental pathways that are essential for the early stage development, and the downstream effector of hippo pathway is YAP/TAZ, which is isolated in drosophila as Yokie. Yokie acted as the transcription activator, and its mutants display the reduced tissue proliferation while overexpression leads to tissue overgrowth. Yap is the mammalian homologue of Yokie, while Taz is the Yap paralog in vertebrates. The function of YAP/TAZ in vertebrates is conserved with the function of Yokie in drosophila: overexpression of non-phosphorylated form of YAP or TAZ increases the proliferation of several human or mouse cell lies in vitro; and depletion of YAP/TAZ by RNAi blocks the growth of several human cancer cell lines. YAP/TAZ are primary sensors of the cell’s physical nature, as defined by cell structure, shape and polarity. YAP/TAZ activation also reflects the cell “social” behavior, including cell adhesion and the mechanical signals that the cell receives from tissue architecture and surrounding extracellular matrix (ECM) [1].

People have found that the cell junction structure of epithelial cells can inhibit the YAP/TAZ activity. YAP/TAZ sense cell polarity and be inhibited by apical signals stimulated by basal signals through phosphorylation of hippo pathway[2]. Studies have shown that disrupting the E-cadherin/a-catenin complex decreased YAP phosphorylation and promoted YAP nuclear accumulation; while disrupting adherin or tight junction turns on the YAP/TAZ nuclear activation[3]. In addition, YAP/TAZ have been shown to be integral components of β-catenin destruction complex that serves as cytoplasmic sink for YAP/TAZ. YAP2 (one of the YAP isoforms) binds to the TJ component ZO-2 (zonula occludens 2) via a PDZ domain, although this interaction appears to favor nuclear translocation rather than membrane retention [4]. In summary, numerous studies have highlighted the interplay betIen the Hippo pathway and apico–basal cell polarity/cell–cell contacts. More and more evidences have shown that Hippo pathway function as a sensor of epithelial architecture/integrity rather than a stereotypical ligand–receptor signal transduction pathway [5]. But there is very limited study to show that YAP/TAZ can regulate the cell junction structure in epithelial cells, except that Yap has been found to be essential to keep cell shape and apical attachment of cortical progenitors [6].

In the nervous system, the neuroepithelial cells are the “stem cells” deriving from actual stem cells in several different stages of neural development[7]. These neural stem cells can differentiate further into multiple types of cells, like neurons, astrocytes and other glial cells. Neuroepithelial cells are a class of stem cell and have similar characteristics, especially the self-renew ability. Before the onset of neurogenesis, neuroepithelial cells (NEs) divide symmetrically to expand their number[8][9]. At the onset of neurogenesis, neuroepithelial cells transform into radial glial cells undergo asymmetric proliferative divisions that give rise to a new neuroepithelial cell and an intermediate progenitor cell. Intermediate progenitor cell will divide symmetrically to generate two neurons. Both neoroepithelial cell and radial glial cell are highly polarized cells, with the apical membrane exposed to the ventricular zone. In the apical membrane, the cell-cell adhesion intermediated by junction proteins, such as beta catenin, E-cadherin, N-cadherin, is an important polarity feature of neuroepithelial cells[7].

Neurons in the brain originate from a relatively small number of neural stem and progenitor cells (NSPCs)[10]. When neurogenesis begins, NEs transform into radial glia (RG) cells that can divide to self-renew and generate a neuron (this is termed direct neurogenesis); alternatively, RG cells can divide to self-renew and generate an intermediate progenitor cell (IPC), which can then divide to generate neurons (this is termed indirect neurogenesis). RG cells can also divide to generate outer radial glia (ORG) cells that can themselves divide to self-renew and generate IPCs or neurons[9].

The VZ has long been thought as a proliferative niche in the developing cortex. HoIver, in contrast with earlier reports of RGPs losing their proliferative capacity and differentiating when they exit the VZ[11]. Asymmetric divisions of radial glia progenitors produce self-renewing radial glia and differentiating cells simultaneously in the ventricular zone (VZ) of the developing neocortex. Whereas differentiating cells leave the VZ to constitute the future neocortex, renewing radial glia progenitors stay in the VZ for subsequent divisions. The differential behavior of progenitors and their differentiating progeny is essential for neocortical development [12].

In this paper, I described YAP/TAZ deletion in mouse neocortex impaired the proliferation and differentiation of neural progenitor cells, and therefore resulted in hydrocephalus. When YAP/TAZ are deleted, both the Sox2 labeled progenitor cell[13] and Tbr2[14] labeled intermediate progenitor cells are decreased in the cortex, meanwhile the ratio of Tbr2/Sox2 is also decreased. The polarity of neuroepithelial cell is also affected when YAP/TAZ are deleted, this is the first time to describe YAP/TAZ regulating polarity proteins in vivo.

## Results

### 1. YAP/TAZ are essential for normal cortex development

To understand the function of YAP/TAZ in the development of nervous system, I deleted Yap and Taz in mouse neocortex through Emx1-cre system. In the YAP/TAZ conditional knockout mice, loss of YAP/TAZ in cortex results in hydrocephalus at the adult stage, the ventricle is significantly dilated. In our investigated total four YapF/F;TazF / Δ;Emx1-Cre mice, all of them displayed the hydrocephalus phenotype; while in the YapF/F;TazF /+;Emx1-Cre mice, only 40% of the display the dilated ventricle phenotype; none of the YapF/+;TazF / Δ;Emx1-Cre mice display the dilated ventricle phenotype. This suggests that YAP/TAZ play essential role in the cortex development, while Yap plays more important role than Taz in the cortex development, and loss of YAP/TAZ leads to hydrocephalus in the mouse.

To determine onset time of the hydrocephalus when YAP/TAZ deleted, I examined the phenotype as early as P0. The result show that conditional deletion of YAP/TAZ in neocortex via Emx1-Cre leads to hydrocephalus as early as P0. Statistic results show that 83% of the mice show the ventricle expansion at P0, this means the most of the hydrocephalus was developed prenatally. The cells linearize the ventricular zone are reduced in the DKO mouse. This indicates the defected mouse cortex may begin at embryotic stage.

To understand the function of YAP/TAZ in the development of nervous system, I detected the expression of YAP/TAZ in the neocortex of mouse from E12.5 to E17.5, the result show that YAP/TAZ are highly expressed and nuclear localized in the progenitor cells, which are marked with Sox2. Meanwhile, in the intermediate progenitor cells, which are marked with Tbr2, I didn’t detect the high expression or nuclear localization of YAP/TAZ.

### 2. Cell polarity features are changed in the DKO neuroepithelial cells

Since Emx1-cre is expressed at Embryotic day 9.5[15], I use IF To determine whether Emx1-cre can deplete YAP/TAZ completely in the neocortex, results show that YAP/TAZ can be completely deleted in the neocortex of mouse at E12.5. I also found that, at E12.5 YAP/TAZ expression are extremely high in the neuroepithelial cells labeled with Sox2. Since neuroepithelial cells are highly polarized, to figure out whether the cell polarity in the ventricular zone are changed when YAP/TAZ are deleted, the polarity marker was used to label the cortex. Result show that the neuroepithelial cell polarity are partially lost in the ventricular zone at E15.5. Since the cells line the ventricular zone are reduced in DKO mice, to figure out whether these cells are progenitor cells I stained the Sox2 positive cells in the P0 cortex. The result show that E-cadherin, N-cadherin and ZO-1 localizations Ire disrupted when YAP/TAZ Ire deleted. The amplified image also show that the beta-catenin is mislocalized in the embryonic day 14.5. The immunofluorescence experiments indicates that the deletion of YAP/TAZ leads to the disrupted neuroepithelial cell junctions.

Polarization is also needed in migrating neuron cells during neurogenesis, postmitotic neurons generated from neuroprogenitor cells will migrate from the ventricle and become paused in a multipolar state within the VZ/SVZ as they extend an axon, the leading edge that faces the direction of migration[7]. Even though this type of polarity is different from apico–basal polarity, it involves many of the same polarity determinants. Since the deletion of YAP/TAZ changed the polarity of neuroepithelial cells, I detected the migration of neural intermediate progenitor cells. The results show that, BrdU labeled intermediate progenitor cells could not migrate to the sub ventricular zone as control in mouse cortex. This indicate YAP/TAZ is required to keep the migration ability of neural intermediate progenitors.

### 3. YAP/TAZ deletion leads to loss of progenitors and intermediate progenitors

Since YAP/TAZ are expressed in Sox2+ cell, to examine the effect of YAP/TAZ deletion on progenitor cells, I analyzed the cell number of Sox2+ cells and intermediate progenitor cells labeled by Tbr2[14]. The statistical analysis results show that, at the apical surface that the ZO-1 is mislocalized, both the Sox2+ and Tbr2+ cells are significantly reduced at E15.5(n=3) (Figure1). The number of Sox2+ cells is reduced when the cell polarity is disrupted in YAP/TAZ deleted mouse cortex, this indicating he progenitor cells are reduced after the deletion of YAP/TAZ. And the ratio of Tbr2/Sox2 is also reduced at the whole apical surface of ventricular zone. Since Tbr2 is a marker for intermediate progenitor cell, and the Tbr2+ cells are born from SOX2 + cells, this indicates that the when YAP/TAZ are deleted, the differentiate ability of SOX2+ cells to Tbr2+ cells is reduced. YAP/TAZ deletion reduced the ability of Sox2+ cells differentiate into Tbr2+ cells.

**Figure1.**
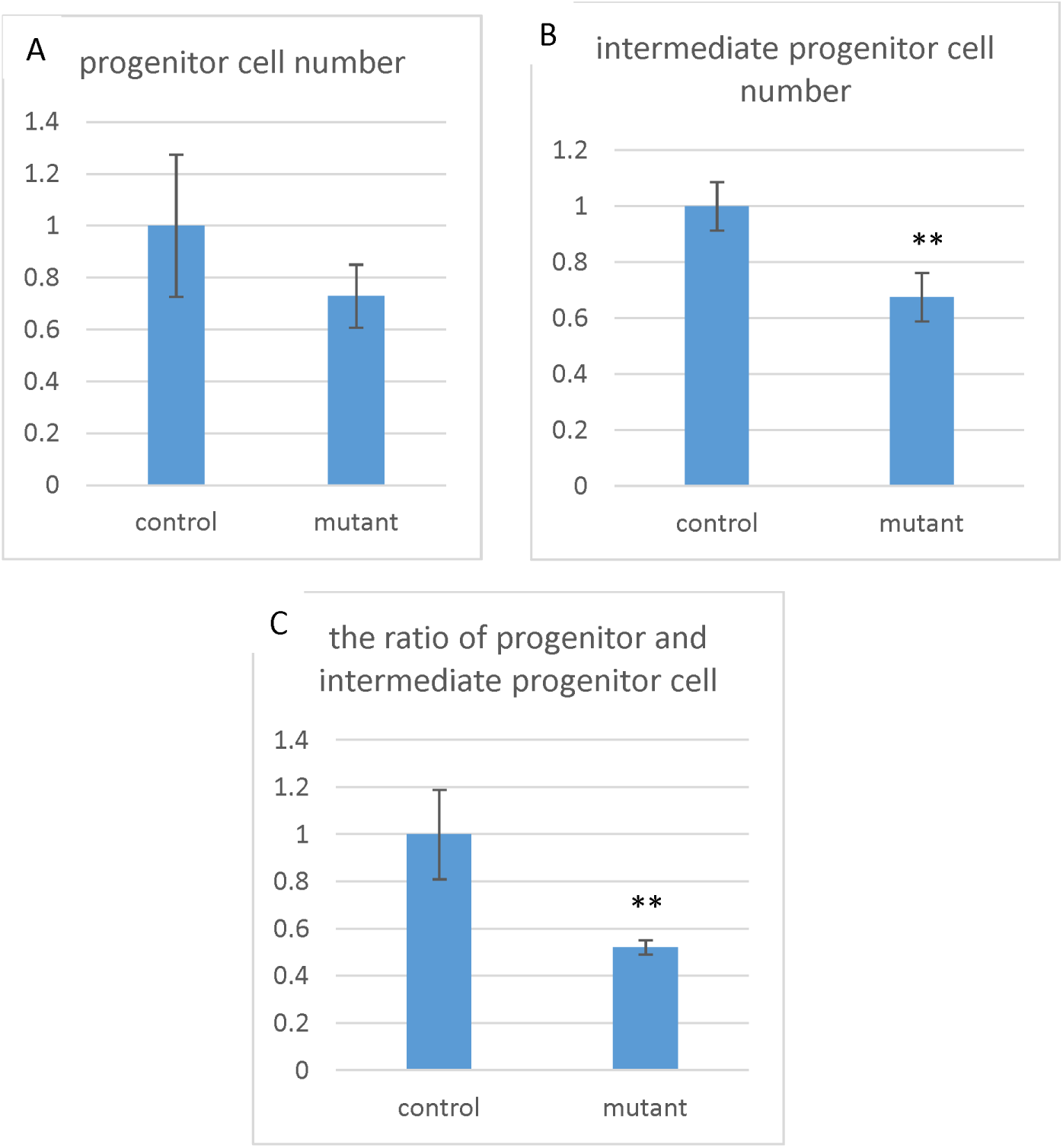
Loss of YAP/TAZ reduced the apical progenitor pool by late neurogenesis in the cortex. (A) SOX2 and TBR2 immunostaining in E15.5 control and YapF/F;TazF/Δ;Emx1-Cre cortex. Scale bar: 20 μm. (B) Quantification of SOX2^+^ and TBR2^+^ cells in E15.5 control and YapF/F;TazF/Δ;Emx1-Cre cortex (*n*□=□3). (C) Quantification of SOX2^+^ (both within the VZ and beyond the VZ) as a percentage of total TBR2^+^ cells in E15.5 control and YapF/F;TazF/Δ;Emx1-Cre cortex (*n*□=□3). Data expressed as mean ± S.E.M. \*\**P*<0.01

### 4. YAP/TAZ deletion increased premature cell cycle exit of neural progenitors

As to neurogenesis, neural progenitors initially divide symmetrically to expand their pool and switch to asymmetric neurogenic divisions at the onset of neurogenesis[10]. To figure out whether the reduced cells are caused by disrupted cell proliferation, I used the EdU labeling to analyze the progenitor cell cycle. I injected EdU into the pregnant mouse at the E14.5, after 24 hours the embryos are collected for analysis. The labeled cells can be detected through immunoflourence with EdU antibody. Ki67 is a cell cycle marker and it is strictly associated with cell proliferating. The proliferation ability of neural progenitor cells can be analyzed through counting the cell number of EdU+Ki67+. The ratio of EdU+Ki67− with EdU+ can display the cell cycle exit ratio. Results show that the YAP/TAZ deletion decreased the proliferation ratio and increased the cell cycle exit ratio, therefore reduced the proliferation ability of progenitor cells. To examine the proliferating index in cell cycle of YAP/TAZ deleted progenitor cells, I co-stained Sox2 and Ki67 in embryonic mouse cortex. The co-staining results of Sox2 and Ki67 show that, in the YAP/TAZ deleted mouse cortex, the double positive cells are reduced compared with control mouse (Figure2). This indicating that more Sox2+ cells exit the cell cycle when YAP/TAZ are deleted, the proliferating ability of neural progenitor cells are impaired. Results of EdU labeling experiment suggested that YAP/TAZ are required to keep the proliferating status of neural progenitor cells, YAP/TAZ deletion increased the cell cycle exit and reduced the proliferation ability of progenitor cells.

**Figure2.**
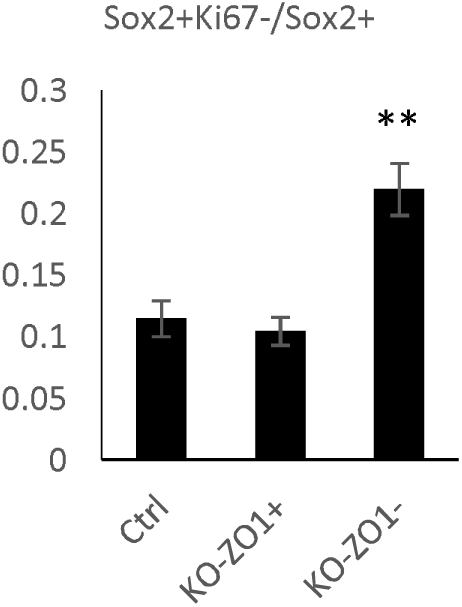
Increased cell cycle exit in the YapF/F;TazF/Δ;Emx1-Cre cortex during early neurogenesis. Pregnant female mice Ire subjected to a 24-hour BrdU pulse prior to sacrifice. Ki67 and BrdU immunostaining in E17.5 control and YapF/F;TazF/Δ;Emx1-Cre cortex. Scale bar: 20 μm. Cell cycle exit indices Ire calculated by measuring the ratio of Sox2+Ki67− cells to total BrdU+ cells in E17.5 control and YapF/F;TazF/Δ;Emx1-Cre cortex (n□=□3). Data expressed as mean ± S.E.M. *P<0.05.

Next I used Tuj1 (Neuron-specific class III beta-tubulin) as a marker to label the differentiated neuron cells, the immunostaining results show that, the TuJ1 is only expressed in sub ventricular zone and barely expressed in ventricular zone. But in the YAP/TAZ deleted mouse cortex, the Tuj1 is highly expressed in the ventricular zone, and localized in the Sox2+ cells. This indicating that, when YAP/TAZ are deleted, the neural progenitor cells express the TuJ1 improperly. The progenitor cells exit the cell cycle and premature into TuJ1+ cells.

## Discussion

In this paper, I demonstrate an essential role for the YAP/TAZ in regulating the proliferation and differentiation of neural progenitors during cerebral cortical neurogenesis in the mouse. Conditional deletion of YAP/TAZ in mouse neocortex via Emx1-Cre leads to hydrocephalus, and this phenotype can be detected as early as P0. In the total four double conditional YAP/TAZ knockout mice, all of them displayed hydrocephalus, 83% (5/6) displayed hydrocephalus at P0. While only 40% of the YapF/F;TazF /+;Emx1-Cre mice display the dilated ventricle phenotype; none of the YapF/+;TazF / Δ;Emx1-Cre mice display the dilated ventricle phenotype. The phenotype of hydrocephalus most likely arises from a loss of progenitors and intermediate progenitors in the ventricular zone of YAP/TAZ deleted cortex. IF results show that YAP/TAZ are expressed in neural progenitor cells and colocalized with Sox2. Both the Sox2+ cells and Tbr2+ cells are significantly reduced at E17.5 when YAP/TAZ are deleted, the ratio of Tbr2/Sox2 is also reduced. This indicates that the differentiation of Sox2+ into Tbr2+ cells is impaired. YAP/TAZ has long been thought to promote proliferation and inhibit differentiation in stem cells[16], but our results display that YAP/TAZ deletion impaired both proliferation and differentiation of neural progenitor cells: The Sox2+ progenitor cells exit cell cycle earlier and premature in to Tuj1+ neurons, the differentiation in to Tbr2+ intermediate progenitors is impaired(Figure3).

**Figure3.**
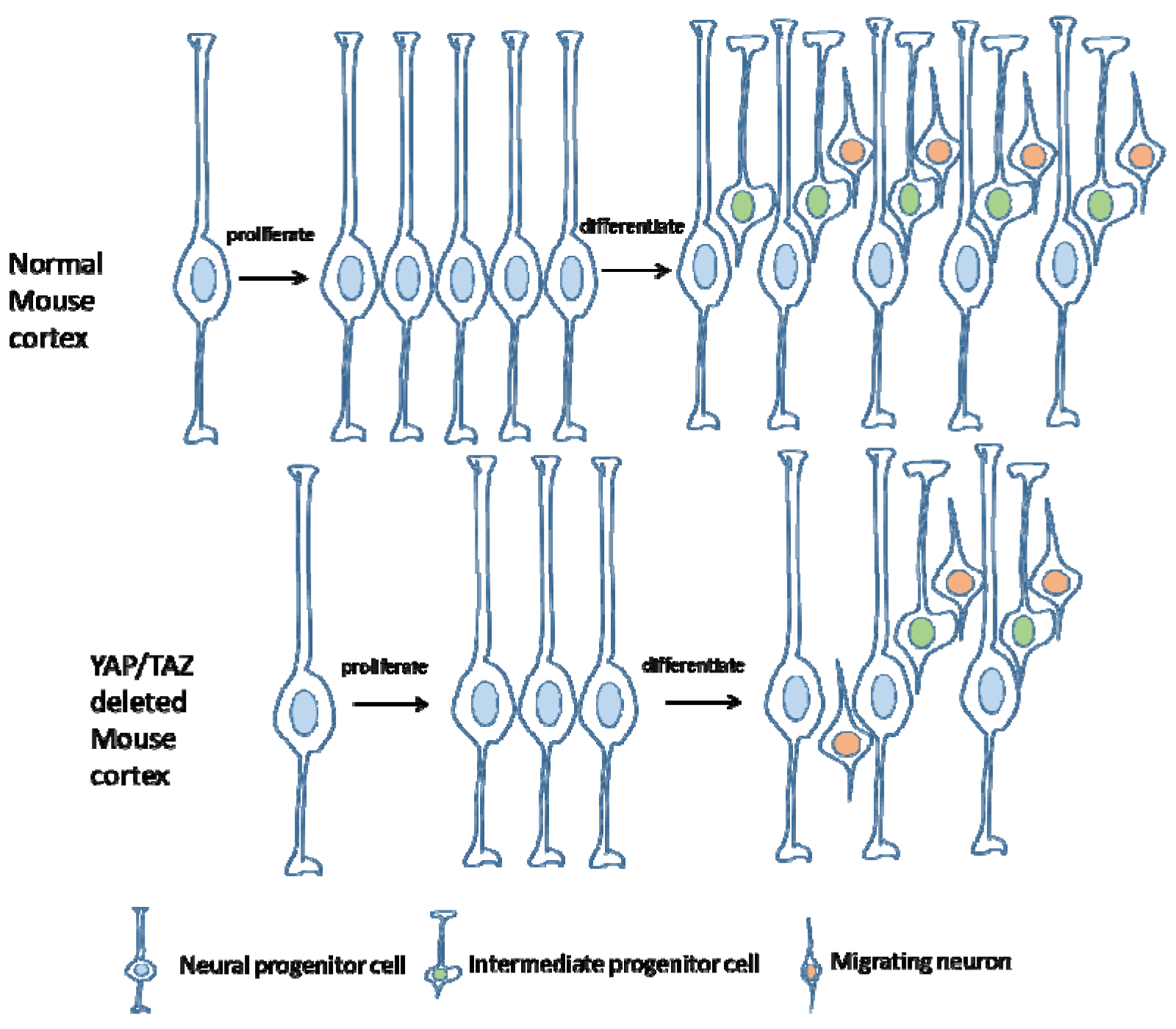
Schematic diagram of the main progenitor cell types and the lineage relationships in the mouse cerebral cortex. Very significant difference exists betIen WT and YAP/TAZ deleted mouse cortex. In normal mouse cortex, the progenitor cell can generate progenitor cell intermediate progenitor cells and neurons through proliferation and differentiation to amplify the pool, the deletion of YAP/TAZ impaired the proliferation and differentiation leading to reduced cell numbers in cortex.

**Figure4.**
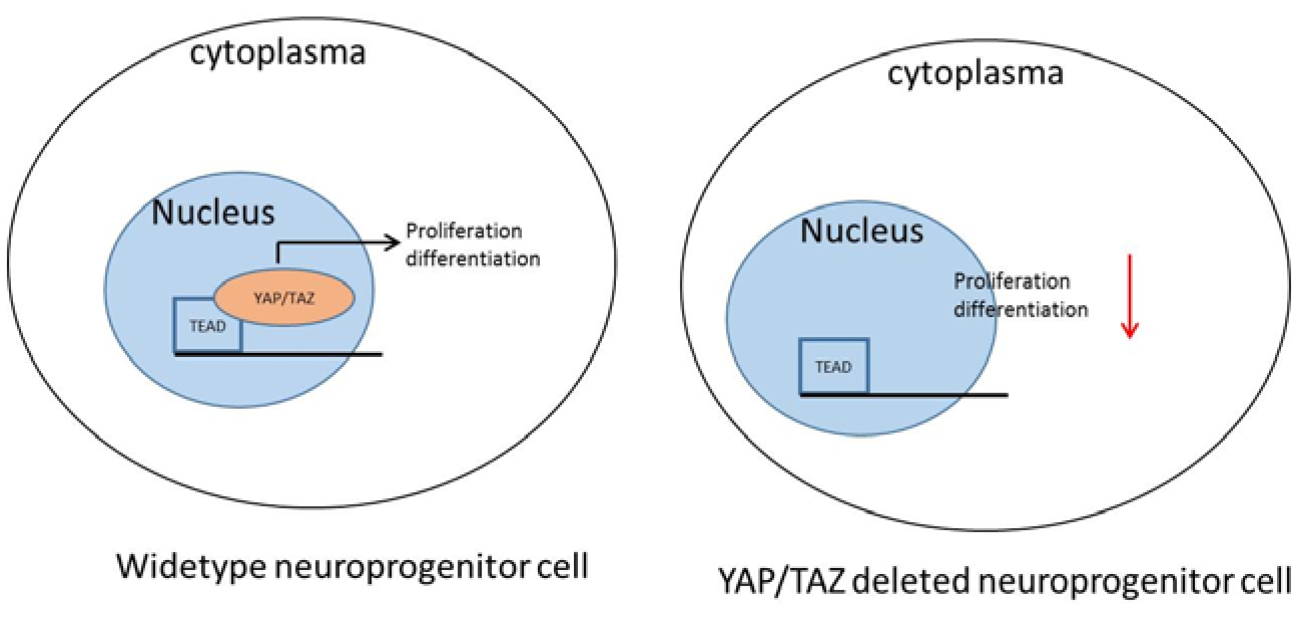
Cartoon pictures illustrate the role of YAP/TAZ in the proliferation and differentiation of neural progenitor cells.

I further analyzed the polarity proteins that are affected by YAP/TAZ deletion and found that, YAP/TAZ can regulate the polarized localization of beta-catenin, and therefore to be essential for keeping the cell polarity. There are several studies to show that YAP/TAZ are regulated by the cell junction and apical-basal polarity of epithelial cells through phosphorylation of hippo pathway[17]. I are the first time to show that YAP/TAZ deletion affected the polarized localization of beta-catenin. This will provide further evidence for the crosstalk betIen Hippo and WNT signaling pathway. Previous studies reported YAP/TAZ as a cell polarity sensor, the polarized cell junction inhibit YAP/TAZ, but in this paper, I found that YAP/TAZ regulate the polarized localization of cell junction proteins, such as beta-catenin and E-cadherin. Our data provide further evidence for the crosstalk betIen YAP/TAZ and Wnt pathway. Taken together, this paper will define the functions of YAP/TAZ in keeping the cell polarity neural progenitors and ensuring their proliferation and differentiation, and also reveal the roles of YAP/TAZ in developing cortex.

YAP/TAZ deletion increased premature cell cycle exit of neural progenitors. Results of EdU labeling and cell cycle analyzing experiments display that YAP/TAZ deletion increased the cell cycle exit, and therefore reduced the proliferation ability of Sox2+ cells. The ratio of EdU+Ki67− cells and EdU+ cells is increased; the Ki67+ progenitor cell ratio is reduced, this indicating that YAP/TAZ deletion increased Sox2+ cell cycle exit, the proliferating ability of neural progenitors is impaired. The improperly increased expression of Tuj1+ in progenitor cells in the YAP/TAZ deleted cortex indicates the premature of Sox2+ progenitor cells. Neuroepithelial cell polarity features are loss. Furthermore, immunofluorescence experiments of polarity proteins show that the deletion of YAP/TAZ disrupted neuroepithelial cell junctions as early as E14.5. Taken together, this paper will define the functions of YAP/TAZ in keeping the cell polarity neural progenitors and ensuring their proliferation and differentiation, and also reveal the roles of YAP/TAZ in developing cortex.

### Mice

Deletion of *YAP/TAZ* in the mouse forebrain was achieved by mating *YAP/TAZ loxP* mice with heterozygous *Emx1Cre* knock-in mice, as previously described. Emx1-Cre (stock no. 005628) was obtained from the Jackson Laboratory.

### EdU labeling

For in vivo labeling of EdU, pregnant mice Ire injected intraperitoneally with EdU at 1 ml per 100 g body Iight (GE Healthcare Life Sciences) and embryos Ire collected and processed for immunofluorescence analysis 24 hours later. To detect EdU positive cells, sections Ire first treated with 1N HCl at 45°C for 30 minutes prior to immunofluorescence analysis. Then washed with TBST, and processed as above using an anti-EdU antibody. For costaining using an anti-EdU antibody and other primary antibodies, sections Ire first stained with the other primary then stained with the anti-EdU antibody.

### Immunostaining

Mouse brains Ire dissected in PBS and fixed overnight in 4% paraformaldahyde in PBS at 4°C. Brains Ire then cryoprotected using 30% sucrose in PBS overnight at 4°C and embedded in OCT for cryosectioning. Frozen sections Ire washed with 0.2% Triton X-100 in TBS (TBST) and incubated in the blocking solution (1% goat serum in TBST) for 1 hour at room temperature. Sections Ire incubated with primary antibodies diluted in the blocking solution overnight at 4°C, washed with TBST, and incubated with secondary antibodies (Jackson ImmunoResearch) diluted at 1:1000 in the blocking solution for 2 hours at room temperature. Sections Ire counterstained with DAPI, washed in TBST, and mounted in ProLong Gold antifade reagent (Invitrogen).

### Antibodies for immunostaining

Primary antibodies: Mouse anti-ZO-1 (Alexa 488conjugated) (Invitrogen 339188); rabbit anti-Phospho-histone H3 (Chemicon 06-570); rabbit anti-Ki67 (Vector Labs VP-RM04); rat anti-BrdU (Abcam ab6326); goat anti-Lhx2 (Santa Cruz sc-19342); goat anti-Sox2 (Santa Cruz sc-17320); rabbit anti-Sox2(Cell Signaling 3728); rabbit anti-Tbr2 (Abcam ab23345); rat anti-Tbr2 (Alexa 647 conjugated, eBiosciences 51-4875); rabbit anti-YAP/TAZ (Cell Signaling 8418 1:500); rabbit anti-YAP (pS127) (Cell Signaling 4911); mouse anti-TUJ1 (1:200; Covance)

Secondary antibodies: donkey anti-rabbit Alexa Fluor 488 (1:500; Invitrogen), donkey anti-goat Dylight 549 (1:1000; Jackson ImmunoResearch), donkey anti-chicken Dylight 649 (1:1000; Jackson ImmunoResearch) and donkey anti-mouse Dylight 549 (1:1000; Jackson ImmunoResearch

### Microscopy and Statistical analysis

Images Ire captured using a Zeiss LSM 510 or 780 confocal microscope. Leica DMI-6000B inverted microscope. For embryonic cortical cell quantifications, 200 μm-wide columns perpendicular to the lateral cortical axis in equivalent regions from three sequential cryosections per brain (*n* □=□ 5) Ire quantified for immunopositive cells. Statistical analysis was performed using Excel 2013 software, and all results are expressed as the mean ± standard error of the mean. *P* values Ire generated using Student’s *t* test (unpaired, two-tailed) to compare betIen two independent data sets and a value of *P<0.05 was considered statistically significant.

